# A High Frequency of Detection of Recombinant Koala Retrovirus (recKoRV) in Victorian Koalas Suggests Historic Integration of KoRV

**DOI:** 10.1101/2024.09.10.612269

**Authors:** Louize Zheng, Alistair R Legione

## Abstract

Recombinant koala retrovirus (recKoRV) is a recently discovered variant of koala retrovirus (KoRV), which likely emerged due to the insertion of another retrovirus (likely Phascolarctos endogenous retrovirus) into the backbone of KoRV. KoRV endogenisation was thought to be ongoing in Victoria based on the low prevalence of the virus based on molecular detection of the *pol* gene, however recKoRV was not incorporated into the previous KoRV diagnostic test results. In this study, a new 5’-region-based PCR assay was developed, capable of detecting both intact KoRV and recKoRV. Using this assay, 319 archived DNA samples from 287 Victorian koalas were retested to investigate KoRV endogenisation. We found a 98.3% (282/287) of these samples were positive for the KoRV-5’ fragment, the majority of which were KoRV-*pol* negative (222/287) on prior testing. Our findings demonstrate extensive KoRV integration into the Victorian koala populations, suggestive of a historic presence of KoRV in Victorian koalas. This finding makes biological sense relative to the translocation history of Victorian koalas, compared to the prior paradigm of ongoing endogenisation, and provides new epidemiological and practical management implications.

## Introduction

Koala retrovirus (KoRV) is a gammaretrovirus that infects koalas (*Phascolarctos cinereus*) and replicates by integrating its proviral DNA into the host genome (Hanger *et al*. 2000). While most retroviruses have inserted into mammalian genomes millions of years ago and are conserved in endogenous, replication-defective forms, KoRV has remained viable in both endogenous and exogenous forms (Tarlinton *et al*. 2006; Hobbs *et al*. 2017). In fact, similarly to other gammaretroviruses that infect cats (feline leukaemia virus) (Rezanka *et al*. 1992), birds (avian leukosis virus) (Neel *et al*. 1981), and apes (gibbon ape leukaemia virus) (Kawakami *et al*. 1980), there is evidence suggesting the clinical correlation of KoRV with leukemia, neoplasia, and chlamydia as secondary diseases (Tarlinton *et al*. 2005; Waugh *et al*. 2017). This makes KoRV an important subject to study for the conservation of koalas as well as for understanding the endogenisation process of retroviruses.

In previous studies, it was found that KoRV can be classified into different subtypes depending on the env sequence, which encodes for the envelope glycoprotein (Chappell *et al*. 2017). KoRV-A is ubiquitously found in all koala populations in Queensland and New South Wales (Waugh *et al*. 2017; Sarker *et al*. 2021) in the endogenous form, that is transmitted vertically (Hobbs *et al*. 2017). In comparison, it was detected in only 20-60% of populations in Victoria and South Australia (Legione *et al*. 2017; Tarlinton *et al*. 2022). KoRV-A in these southern populations was suspected to be not fully endogenised, and could also present the exogenous form to be transmitted horizontally (Wedrowicz *et al*. 2016). Based on the epidemiological distribution, KoRV-A is considered to have originally spread from north to south, with its endogenisation process currently taking place in the same direction (Tarlinton *et al*. 2006; Simmons *et al*. 2012; Blyton *et al*. 2022). Moreover, as identified by other studies, there are multiple subtypes (Shimode *et al*. 2014; Chappell *et al*. 2017), which have likely emerged from mutations of KoRV-A within infected koalas (Blyton *et al*. 2022). These are observed exclusively in the exogenous forms, and they show great prevalence variabilities within each population (Legione *et al*. 2017; Waugh *et al*. 2017). These include a more pathogenic subtype, KoRV-B, that is found more frequently in the northern populations, similarly to KoRV-A (Legione *et al*. 2017; Waugh *et al*. 2017; Quigley *et al*. 2018). Hence, such geographical patterns of KoRV epidemiology can indicate susceptibility of a chosen koala population to secondary diseases.

However, in recent studies, long-read DNA sequencing has led to the discovery of a disrupted form of KoRV. This new variant is called recombinant KoRV (recKoRV), in which the mid-region of KoRV is substituted with Phascolarctos endogenous retroelement (PhER) as shown in Figure 1 (Löber *et al*. 2018; Tarlinton *et al*. 2022). As a result, it has lost the entire *pro-pol* gene, some *gag* and some *env* regions, that were critical for the virus detection in PCR based diagnostics (Tarlinton *et al*. 2005; Tarlinton *et al*. 2022). This has raised a question whether previously identified KoRV-negative koalas are in fact naïve to an endogenous form of this virus. The presence of recKoRV in southern koalas could indicate KoRV has already entered their ancestral germlines, and the endogenisation process is more advanced than previously believed. Additionally, it may suggest recKoRV is somehow providing protection against KoRV-A and KoRV-B infection in southern populations. Therefore, to determine the recKoRV prevalence in Victorian koalas, we used 319 archived samples previously tested for the KoRV *pol* region, to run PCR targeting an alternative, intergenic region conserved in both intact KoRV and recKoRV.

**Figure 1.**
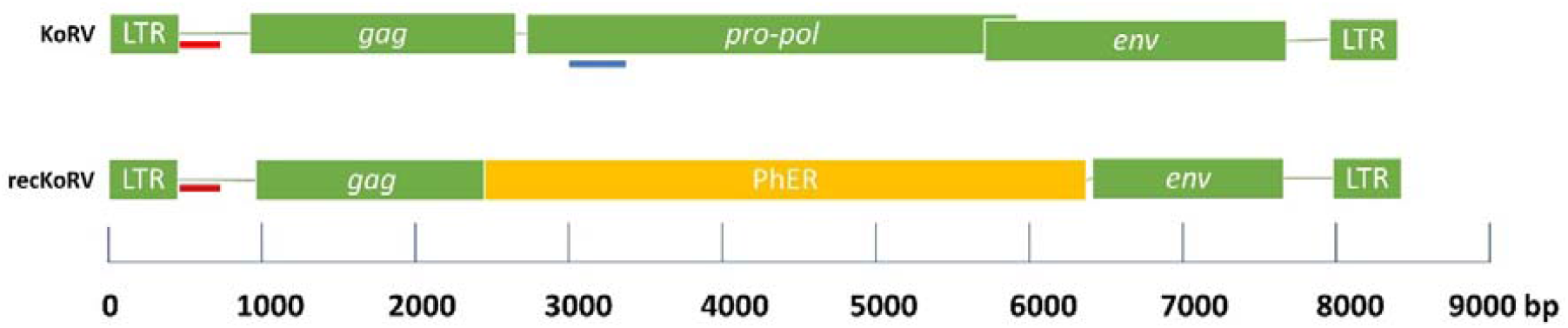
Genomic structures of koala retrovirus (KoRV) and recombinant koala retrovirus (recKoRV). Red bars represent the 5’ region amplified by the primers in this assay. A blue bar represents the target region in previous pol-based PCR assays. LTR: long terminal repeat. PhER: Phascolarctos endogenous retroelement. bp: base pairs.

## Methods

### Sample Collection

A total of 319 archived samples from 300 Victorian koalas were used in this research. These samples, collected between 2010 and 2015, were previously tested for the presence of KoRV provirus (Legione *et al*. 2017). DNA was extracted using the Corbett Xtractor robot and Qiaxtractor VX extraction kit as per manufacturer’s instructions and had been stored in 96-well extraction plates at −20 °C prior to the recKoRV testing. The sample types targeted were buffy coat and spleen samples, with incidental inclusion of whole blood, plasma, and serum sample extracts if they were present in the original extraction plates. Each sample was associated with previous records of location, sex, age, β-actin genomic copy number per extraction, and KoRV provirus copy number per extraction based on *pol*-targeted qPCR.

Additionally, initial testing utilised recently obtained diagnostic samples from koalas from Queensland and Victoria. These samples included skin, blood, and spleen samples that were extracted using the Wizard® Genomic DNA Purification Kit (Promega). Three samples from Victorian koalas that had previously been identified as containing recKoRV via long read sequencing were also included in initial testing (Tarlinton *et al*. 2022).

### Primer Design

Two sets of primers (Table 1) were designed using primer3plus 2.3.7 (Untergasser *et al*. 2007) in Geneious Prime 2023.01 (Kearse *et al*. 2012). Both were developed to target a 5’ intergenic region found in KoRV and recKoRV (Figure 1), based on the Biosample sequences of SAMN23247354, SAMN23247355, SAMN23247356, and SAMN23247357 (Tarlinton *et al*. 2022) aligned against KoRV-A (NC039228) (Hanger *et al*. 2000). Briefly, the FASTQ reads generated in that study were assembled using Flye (Kolmogorov *et al*. 2019), and aligned using MAFFT (Katoh and Standley 2013), to determine conserved sites across multiple recKoRVs (FASTA sequences available in Supplementary File 1). In our study, samples successfully amplified with the primers were defined to be “KoRV-5’ positives”. They were distinguished from “recKoRV positives”, which had more copies of the 5’ fragment than the *pol* region based on qPCR absolute quantification.

**Table 1.**
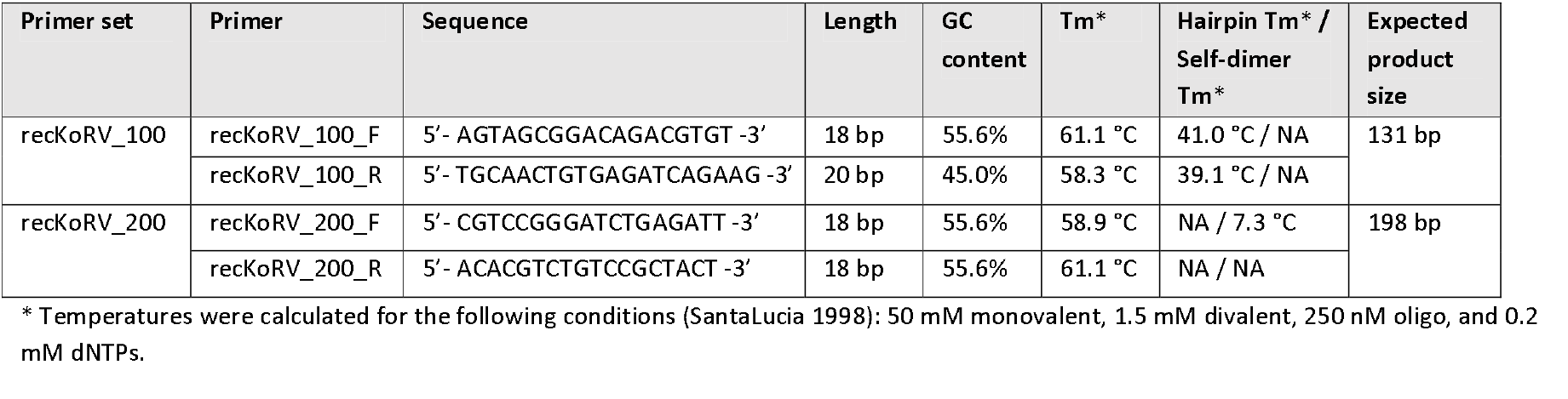
Primer designs for KoRV and recKoRV detection in PCR.

### Initial Testing of Primers

Conventional PCR was utilised with three diagnostic samples of unknown recKoRV status and three known recKoRV positive samples (all six were previously tested on *pol*-PCR and contained three KoRV-negatives and three KoRV-positives). This was performed in a T100 Thermal Cycler (Bio-Rad). Each reaction mixture consisted of 5 µL template, 500 nM of each primer, 0.20 mM of each dNTP, 2.00 mM MgCl_2_, 1.5 units GoTaq Flexi DNA Polymerase (Promega), and 1 × GoTaq Flexi buffer. The final volume was made up to 25 µL using nuclease free water. The thermal profile of the PCR was as follows: initial denaturation at 95 °C (four minutes), 40 cycles of denaturation at 95 °C (30 seconds), annealing at 55 °C (30 seconds), and extension at 72 °C (15 seconds). The final cycle was followed by a final extension at 72 °C for five minutes. Consequently, the products were visualised using a 1% w/v agarose gel electrophoresis, in conjunction with HyperLadder 100 bp (Bioline) to determine the presence of the target sequences. To confirm their specificities in target amplification, four PCR products (SAMN23247356 and SAMN23247357 amplified with each of the two primer sets) from the following section were sequenced using Sanger dideoxy sequencing at the Australian Genomic Research Facility and aligned against the original KoRV-A reference sequence (RefSeq Accession: NC039228).

### Plasmid Control Preparation

To create positive controls and standards for quantitative PCR (qPCR), a known recKoRV-positive sample (SAMN23247356) was amplified in the conventional PCR assay with primers recKoRV_100 and recKoRV_200. These PCR products were purified with the QIAquick Gel Extraction Kit (Qiagen) and then ligated into pGEM-T Easy plasmids (Promega) overnight. Ligated plasmids were transformed into NEB 5-alpha competent E. coli (New England BioLabs) using manufacturer’s protocol and plated onto Luria-Bertani (LB) agar plates with ampicillin for antibiotic selection, IPTG and X-gal for blue/white selection. Selected colonies were cultured overnight at 37°C in LB broth in the presence of ampicillin and shaking at 200 rpm. The plasmids were extracted from 5 mL of overnight culture using the Wizard® Plus SV Miniprep DNA Purification System (Promega), and the concentration of purified plasmid was determined using a Qubit 3.0 Fluorometer (Invitrogen). Genomic copy numbers were calculated from this value, and for each plasmid, 10-fold dilutions from 10^8^ to 10^1^ copies per 5 µL were prepared, in nuclease free water, with a QIAgility robot (Qiagen).

### PCR Optimisation

Positive plasmid controls (10^1^, 10^2^, 10^3^, 10^4^, 10^5^, 10^6^, 10^7^, 10^8^ copies per 5 µL) and a non-template control (nuclease free water) were used to undertake optimisation of annealing temperature (51-62 °C), primer set (recKoRV_100, recKoRV_200), MgCl_2_ (1.5 and 2.0 mM), and primer concentration (recKoRV_100: 0.1, 0.2, 0.3, 0.4, 0.5 µM; recKoRV_200: 0.25 and 0.5 µM). These were carried out with reagents in Table 2 and the basic thermal cycling conditions as described previously. Conventional PCR was used for temperature optimisation, and qPCR for the remaining tests. For qPCR, SYTO™ 9 Green Fluorescent Nucleic Acid Stain (ThermoFisher Scientific) was incorporated into the master mix at a concentration of 2.0 µM per reaction.

**Table 2.** Reagents and thermal setup in PCR optimisation.

|  | Primers |  | MgCl<sub>2</sub> | Annealing temperatures | Dilution range tested (copies / reaction) |
| --- | --- | --- | --- | --- | --- |
|  | Set | Concentration |  |  |  |
| Annealing temperature | recKoRV_100 | 0.50 µM | 2.00 mM | 53, 55, 57, 60, 62 °C | 10<sup>1</sup>, 10<sup>2</sup>, 10<sup>3</sup>, 10<sup>4</sup> |
|  | recKoRV_200 |  | 1.50 mM | 51, 53, 55, 58, 60 °C | 10<sup>1</sup>, 10<sup>2</sup>, 10<sup>3</sup> |
| MgCl<sub>2</sub> concentration | recKoRV_200 | 0.50 µM | 1.50 mM | 55 °C | 10<sup>1</sup>, 10<sup>2</sup>, 10<sup>3</sup>, 10<sup>4</sup>, 10<sup>5</sup>, 10<sup>6</sup>, 10<sup>7</sup>, 10<sup>8</sup> |
|  |  |  | 2.00 mM |  |  |
| Primer set | recKoRV_100 | 0.50 μM | 1.50 mM | 55 $^{\circ}$C | 10<sup>1</sup>, 10<sup>2</sup>, 10<sup>3</sup>, 10<sup>4</sup>, 10<sup>5</sup>,<br>10<sup>6</sup>, 10<sup>7</sup>, 10<sup>8</sup> |
|  | recKoRV_200 |  |  |  |  |
| Primer concentration | recKoRV_100 | 0.1, 0.2, 0.3, 0.4, 0.5 μM | 2.00 mM | 55 $^{\circ}$C | 10<sup>1</sup>, 10<sup>2</sup>, 10<sup>3</sup>, 10<sup>4</sup>, 10<sup>5</sup>,<br>10<sup>6</sup>, 10<sup>7</sup> |
| | recKoRV_200 | 0.25, 0.50 μM | 1.50 mM | | |

### Detection of Endogenous KoRV-5’

The archived samples of DNA extracts were tested for endogenous KoRV-5’ using an AriaMx real-time PCR system (Agilent) with recKoRV_200 primers under the optimised conditions (see results). A melt curve was obtained at a 0.5 °C resolution from 65 °C to 95 °C in order to confirm the target amplification. The copy numbers in the original samples were estimated against the standard curve, acquired from the positive controls of 10-fold dilutions (10^8^-10^2^ copies per reaction) in triplicate. A sample was reported as “KoRV-5’ positive” if it met both of the following conditions: 1) the detected copy number was above 100 per reaction, and 2) the melt curve had a dominant peak in the range of 84 to 86°C. Potential positives with a melt curve peak between 83 and 87 °C, or with a copy number detected below the threshold, were assessed in gel electrophoresis to determine the KoRV-5’ positives.

### Statistical Analyses

For our samples, qPCR absolute quantification was used to determine KoRV-5’ genomic target copy numbers per reaction, and compared to previously obtained KoRV-*pol* and β-actin genomic copy numbers (Legione *et al*. 2017). A crude estimate of recKoRV genomic copy numbers were obtained by subtracting the previous KoRV-*pol* copy numbers from our KoRV-5’ copy numbers. For the KoRV-5’ prevalence, univariable regression model was used to investigate any statistical correlation of the results with the demographic (sex, age, and region) and contextual (sample type and extraction plate) variables. Non-parametric statistical tests were performed with the software ggstatplot (Patil 2021), as copy numbers were not normally distributed even after logarithmic conversion. β-actin genomic copy numbers were compared using a Kruskall-Wallis test, by sample type and by their results in *pol*- and 5’-PCR assays. Mann-Whitney U test was used to compare the KoRV-5’ and the estimated recKoRV genomic copies between KoRV-*pol* positive and negative samples. Spearman’s Rank-Order Correlation was then performed to examine the correlation of recKoRV and KoRV-*pol* genomic copies.

## Results

### Primer Specificities in the Initial Testing

All six koala samples tested positive for KoRV-5’ (Figure 2). Two negative control samples (a non-host control using poultry cells, and a nuclease free water control) showed no target amplification. Using BLAST (Altschul *et al*. 1990), PCR products of SAMN23247356 and SAMN23247357 both showed 99.0% (for reckKoRV_100) and 99.4% (for recKoRV_200) nucleotide identities with XM_020970715.1 (PREDICTED: *Phascolarctos cinereus* uncharacterised LOC110197110, mRNA), which when analysed in this study was an integrated recKoRV in the koala reference genome.

**Figure 2.**
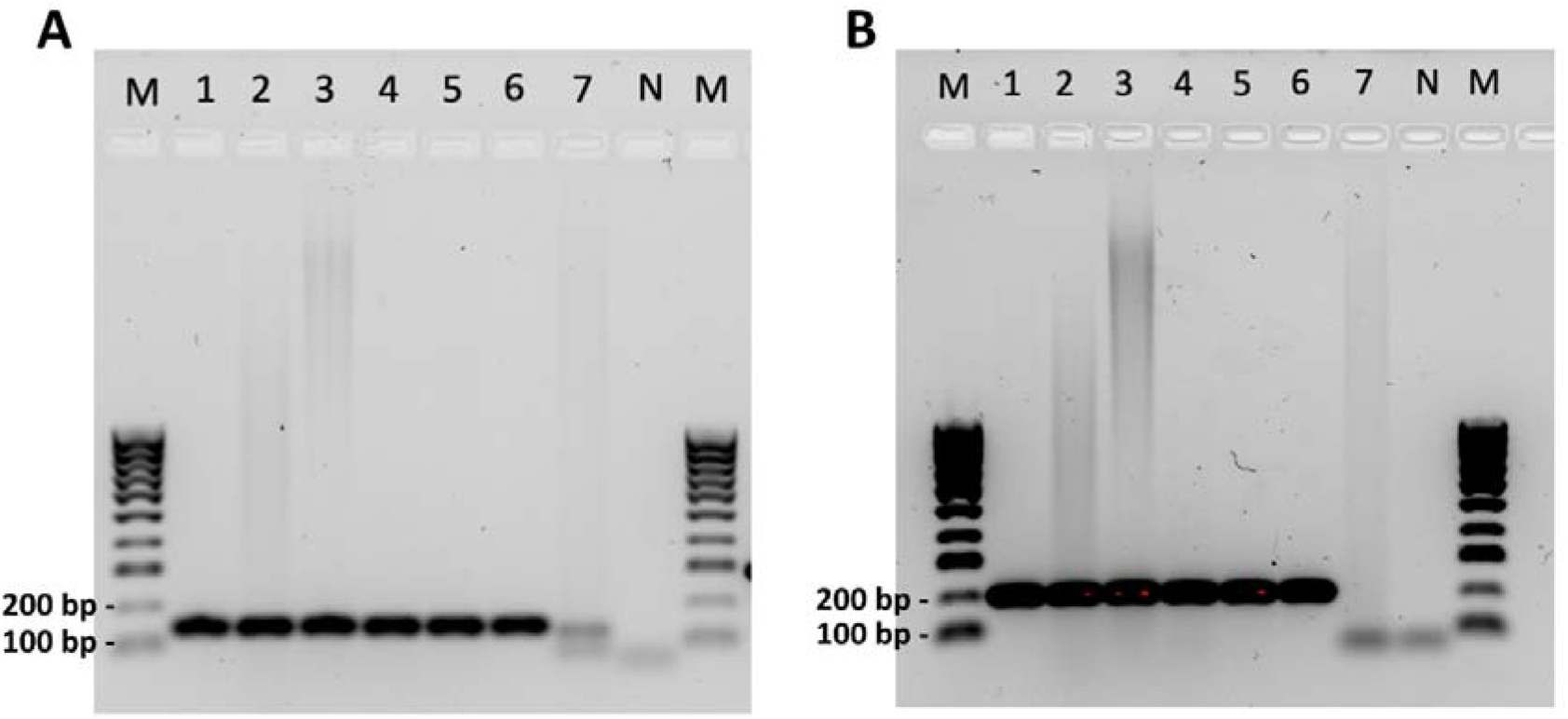
Gel images of the initial primer testing results for (a) recKoRV_100 and (b) recKoRV_200. M: HyperLadder 100 bp (Bioline). 1: diagnostic skin sample from Queensland (KoRV positive). 2: diagnostic spleen sample from Victoria (KoRV negative). 3: diagnostic blood sample from Victoria (KoRV positive). 4: SAMN23247355 (KoRV positive, known recKoRV positive). 5: SAMN23247356 (KoRV negative, known recKoRV positive). 6: SAMN23247357 (KoRV negative, known recKoRV positive). 7: non-host control. N: negative control.

### Optimised PCR Conditions

For primer sets, the minimum qPCR detection limit was 10^3^ copies per reaction for recKoRV_100, and 10^1^ copies per reaction for recKoRV_200. A higher specificity was reached for recKoRV_200, for which dilutions of at least 10^2^ copies per reaction showed a single peak at target temperatures (84-86 °C) in melt curve analysis (MCA). In contrast, recKoRV_100 showed off-target MCA peaks in all dilutions. In the conventional PCR, the optimal annealing temperature was 55°C for both recKoRV_100 and recKoRV_200, based on the strength of the visualised bands. However, off-target products, likely dimers (< 100 bp), were observed at all temperatures and for all dilutions. In qPCR, 1.50 mM MgCl_2_ showed a dominant target MCA peak for dilutions of at least 10^1^ copies per reaction, whilst 2.00 mM MgCl_2_ could not detect a target MCA peak in the 10^1^ copy per reaction. For recKoRV_100, 0.2, 0.3, and 0.5 µM primer concentrations all had the minimum detection limit of 10^1^ copies per reaction in qPCR, although they created off-target MCA peaks in all dilutions. For recKoRV_200, 0.5 µM had the same minimum detection limit, but off-target peaks were no longer present in 10^7^- and 10^8^-copy dilutions. At 0.25 µM, recKoRV_200 had a slightly worse minimum detection limit of 10^2^ copies per reaction, but the detected dilutions did not have any major off-target peaks. This final condition showed no dimers for dilutions down to 10^2^ copies per reaction when assessed on gel.

Based on the optimisation results, all subsequent qPCRs were run using recKoRV_200 primers, with 250 nM of each primer, 1.5 mM MgCl_2_, 200 nM of each dNTP, 2 µM of SYTO9, 1.5 Units of GoTaq *pol*ymerase (Promega) and 1 × GoTaq Flexi Buffer. All PCR reactions were made up to 20 µL with water followed by the addition of 5 µL of template. The cycling protocol was 95 °C for 4 minutes, followed by 40 cycles of 95 °C for 30 seconds, 55 °C for 30 seconds, and 72 °C for 15 seconds. A final extension at 72 °C for 5 minutes was undertaken, followed by melt curve generation as previously described.

### KoRV-5’ Prevalence

The overall KoRV-5’ prevalence across Victorian koala samples was 98.3% (95% confidence interval (CI): 96-99%, 282/287), with each region (as previously described in Legione *et al*. (2017)) having KoRV-5’ detection between 97 and 100% of samples tested (Figure 3). The data are broken down by demographic and contextual variables in Supplementary Table 1. No variables were significantly associated with the KoRV-5’ detection.

**Figure 3.**
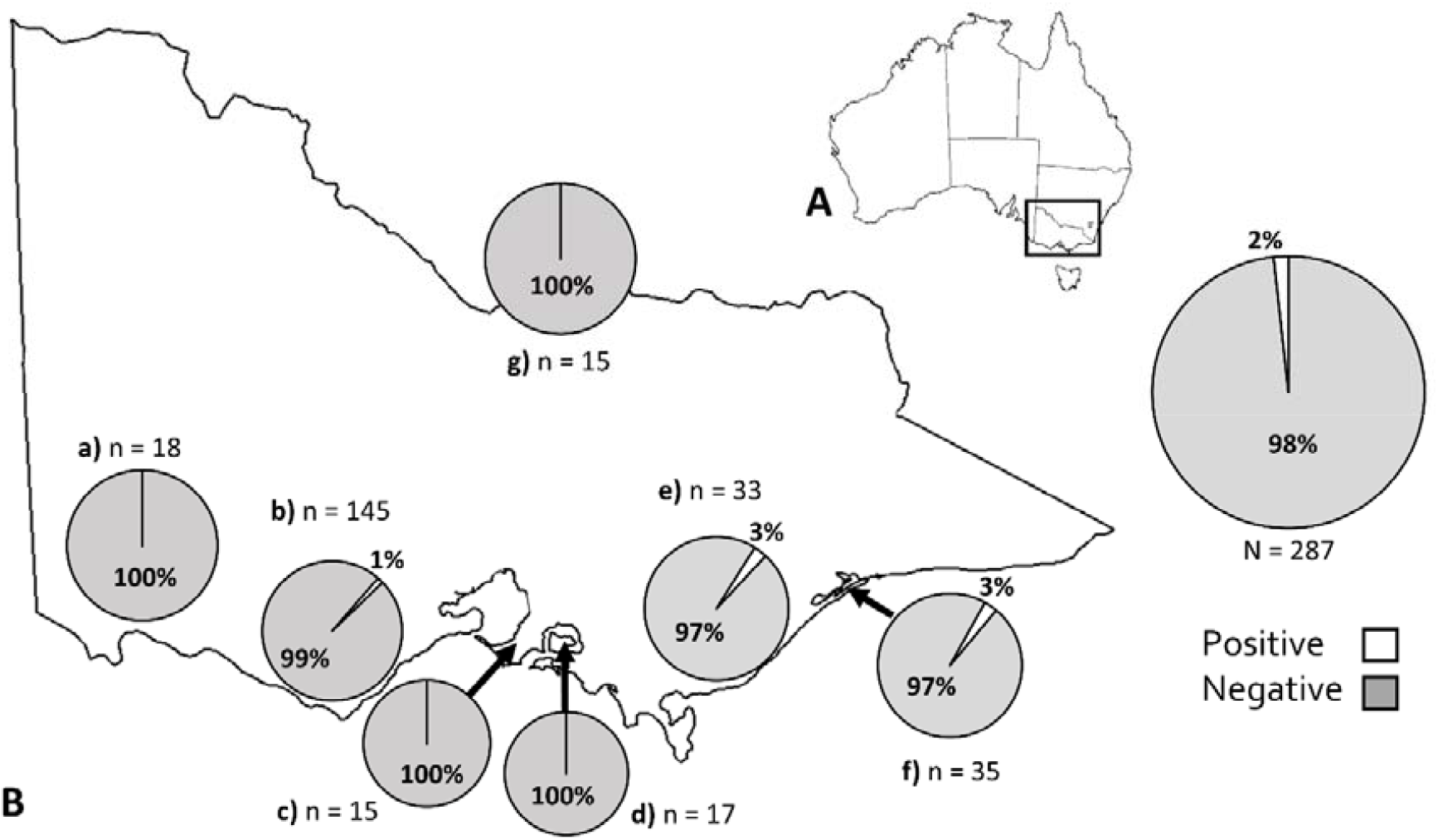
KoRV-5’ prevalence in Victorian koala populations. (A) Location of Victoria in Australia. (B) KoRV-5’ prevalence in individual koala populations which had more than 10 animals tested: (a) Far West, (b) South Coast, (c) Mornington Peninsula, (d) French Island, (e) Gippsland, (f) Raymond Island, and (g) Far North.

Within each sample type, the β-actin genomic copy numbers recorded in a prior study were found to be comparable in KoRV-5’ positive cases regardless of KoRV-*pol* status (Kruskall-Wallis test; buffy: p = 0.148, spleen: p = 0.76, whole blood: p = 0.67). Between sample types, there was a significant difference between the β-actin genomic copy numbers (Figure 4), and thus all further comparisons were based on a ratio of test genomic copies (KoRV-5’ or KoRV-*pol*) per β-actin copies detected.

**Figure 4.**
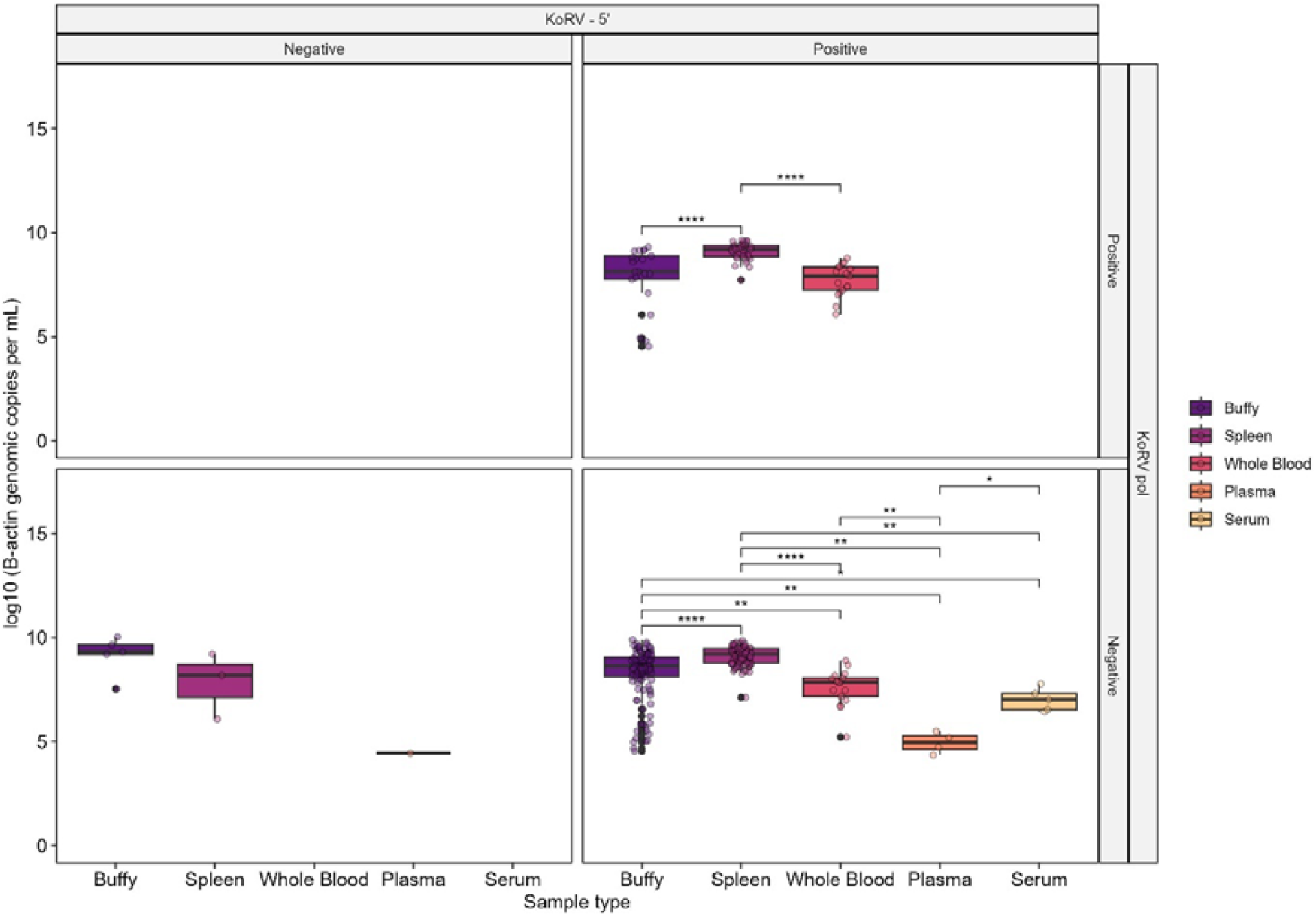
Comparison of β-actin genomic copy numbers in samples with different results of the KoRV-*pol* and KoRV-5’ assays. (*: p ≤ 0.05, **: p ≤ 0.01, *** p ≤ 0.001, **** p ≤ 0.0001)

### Detection of Two Different KoRV Genome Fragments

Of the 287 koalas tested across Victoria, a majority were KoRV-*pol*-negative and KoRV-5’-positive (217/287, 75.6%, 95% CI: 70.2-80.5%). This was followed by animals positive for both KoRV fragments (65/287, 22.7%, 95% CI: 17.9-27.9%) and animals negative for both KoRV fragments (5/287, 1.7%, 95% CI: 0.57-4.0%). No cases were KoRV-*pol*-positive and KoRV-5’-negative.

The median ratio of estimated recKoRV copies per β-actin was 0.27 (1st quartile: 0.155, 3rd quartile: 0.070). The median KoRV-5’ copy numbers per β-actin was statistically higher in KoRV positive animals compared to KoRV negative animals (Mann-Whitney U test, p = 0.04), however when removing KoRV-*pol* copies from the KoRV-5’ values and only looking at estimated recKoRV copy numbers, this significance was removed (Figure 5). In KoRV-*pol* positive, recKoRV positive animals, a majority (41/54, 75.9%) had more copies of recKoRV than KoRV-*pol* while the recKoRV copy in each individual was at a comparable level regardless of the KoRV-*pol* abundance (Spearman = −0.01, p = 0.92)(Figure 6).

**Figure 5.**
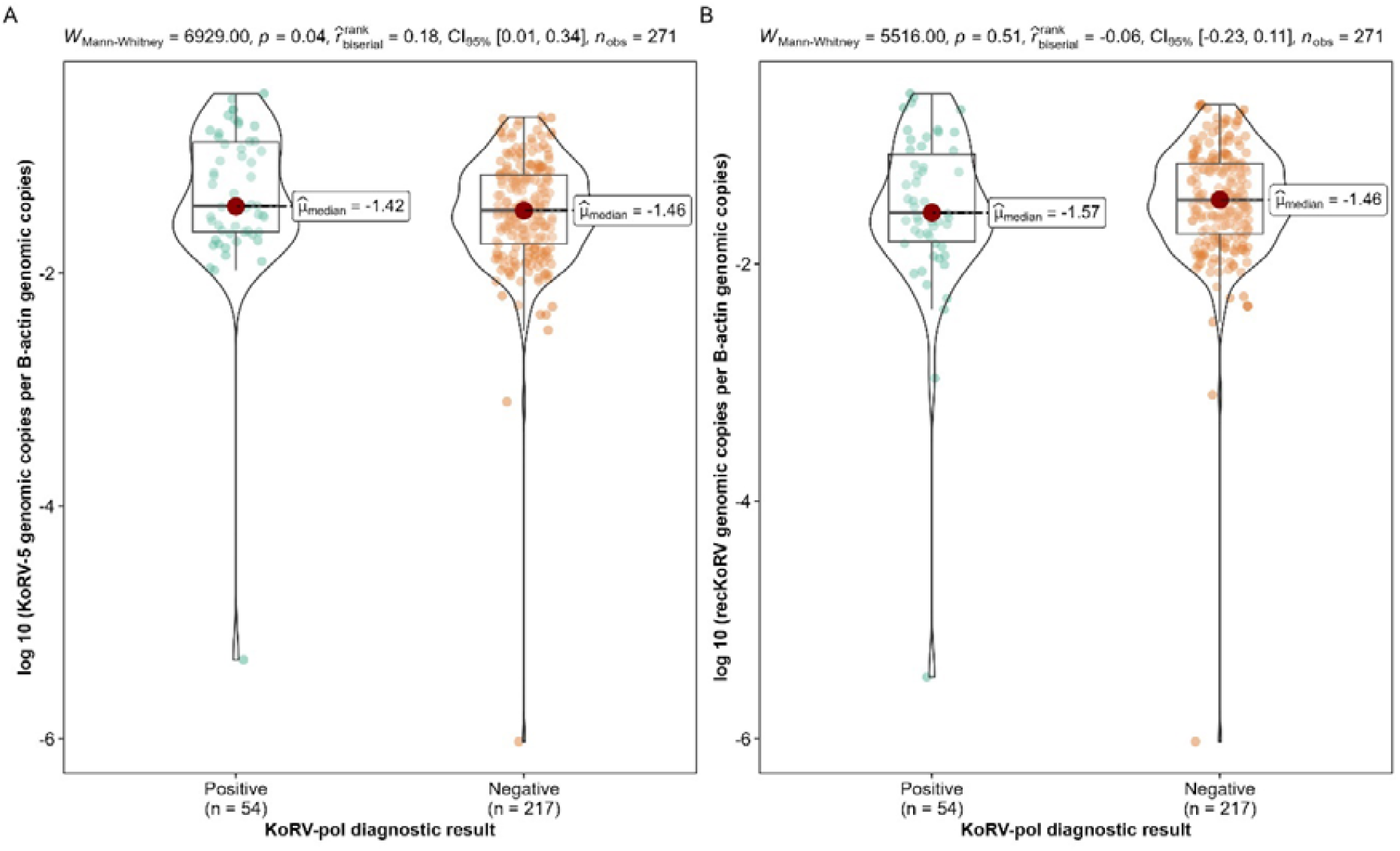
Comparison of the (A) KoRV-5’ and (B) estimate recKoRV genomic copy numbers by KoRV-*pol* status.

**Figure 6.**
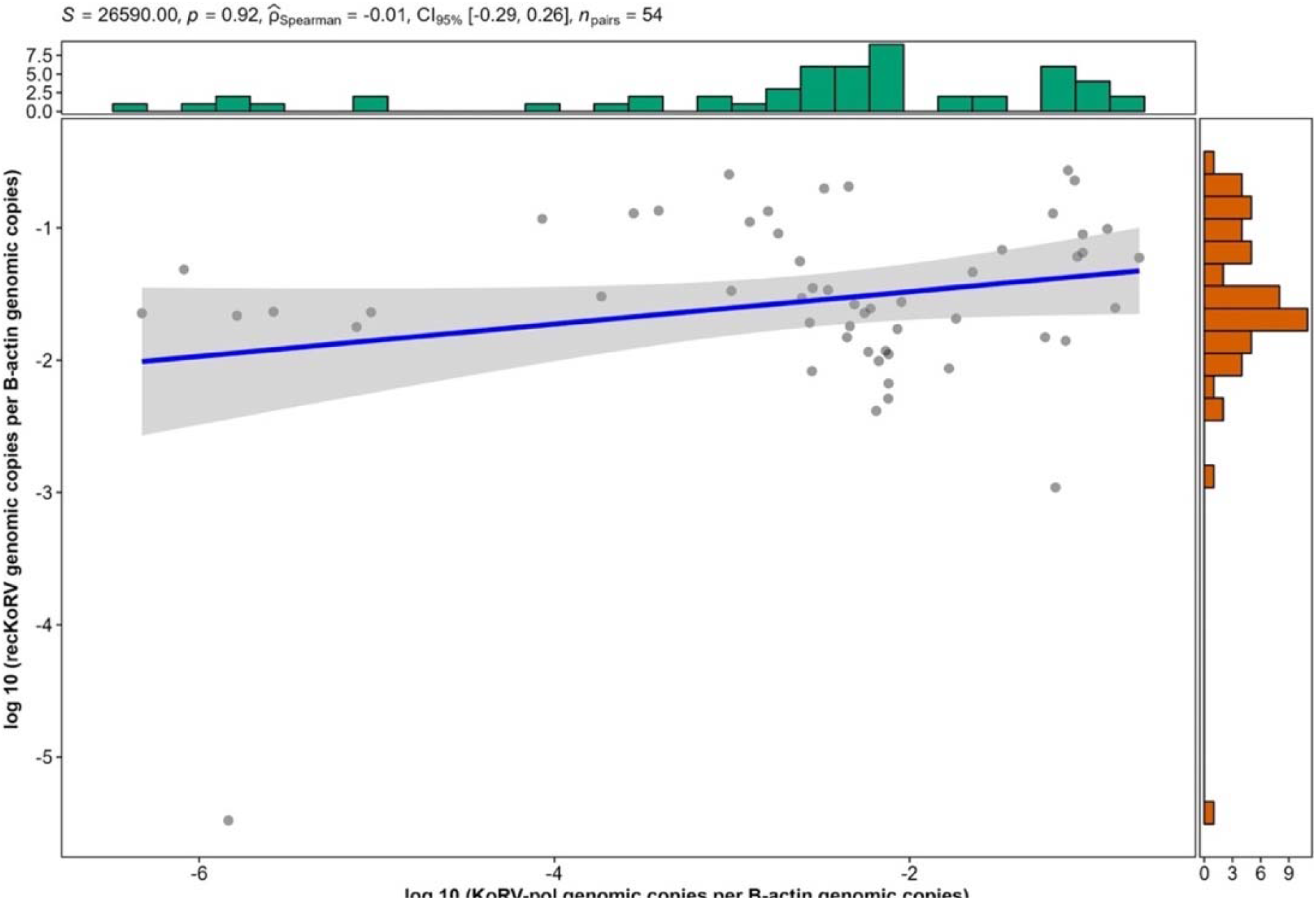
Correlation of the genomic copy numbers of recKoRV and KoRV-*pol* in individual animals

## Discussion

In this study, the KoRV-5’ fragment was detected in all Victorian koala populations. This is a significant finding that contrasts with the 25% KoRV-*pol* prevalence reported previously using the same samples but an alternate target region (Legione *et al*. 2017). While this assay was not able to distinguish between intact KoRV and recKoRV due to the target site used, the stark difference in the numbers indicates a likely high prevalence of recKoRV among Victorian koala populations. This was previously suspected from the genome sequencing of four southern koalas (Tarlinton *et al*. 2022), three of which were from Victoria. By testing Victorian koala populations at a much larger scale our study has provided strong supporting evidence with the high recKoRV prevalence suggesting that KoRV has largely been endogenised in Victorian koalas. In particular, the high KoRV-5’ prevalence in French Island may explain why the KoRV prevalence had seemingly flattened at 25% in the small, closed population (Legione *et al*. 2017). Given this is one of the oldest remaining populations in Victoria (Menkhorst 2008), it appears the KoRV endogenisation has occurred at a much earlier time point (before koala’s were introduction to French Island in the 1890s (Lewis 1934)) than what was previously expected based on the apparent low KoRV prevalence of Victorian koalas (Tarlinton *et al*. 2006; Simmons *et al*. 2012; Legione *et al*. 2017).

Presumptively intact KoRV (determined by KoRV-5’ positive and *pol* positive) was present in less than a quarter of the KoRV-5’ positive animals we assessed in this study. This result highlights that recKoRV is the dominant endogenised form of the virus in Victorian koala populations. A study in South Australia by Stephenson *et al*. (2021) found similar results to our study, with a 99.5% detection rate of a 5’ gag region in koalas despite only obtaining a 41% positive result for KoRV-*pol*. That same gag region is present in the assembled contigs that we generated from the sequence confirmed recKoRV positive animals previously described (Tarlinton *et al*. 2022). South Australian koalas have historical origins in Victoria, with many populations originally derived from translocations from French Island in Victoria (along with other Victorian populations) (Menkhorst 2008), it is likely the koalas in both Victoria and South Australia have similar endogenisation of recKoRV.

At an individual level, where animals in Victorian populations had both the intact and broken forms of KoRV, the majority had more recKoRV than full-length KoRV genomic copies based on comparison of qPCR results. This is in contrast to the observation made in northern koalas, where the recKoRV and KoRV integration sites were found in a 1:3 (Löber *et al*. 2018) or 1:4 (Hobbs *et al*. 2017) ratio. The higher KoRV disruption rate in Victorian populations further emphasises the discrepancy in their true and apparent levels of KoRV endogenisation. Nevertheless, the total KoRV genomic copy numbers (including intact KoRV and recKoRV) were previously found at a significantly higher level in northern koalas than southern koalas (Simmons *et al*. 2012; Blyton *et al*. 2022). This remains true even when the results for Victorian koalas are adjusted to include the recKoRV copy numbers in our assay. It should be noted that because we utilised old extractions, our samples may contain less DNA, due to degradation, than what the previously obtained β-actin numbers indicate (Legione *et al*. 2017). Whilst no samples returned KoRV-5’-negative, several samples had lower KoRV-5’ copies than KoRV-*pol* copies. It is most likely these samples are representative of minor degradation, as the only alternate hypothesis is that there is an undescribed integrated fragment that lacks the targeted 5’ region, but contains an intact *pol* target region. Our results contrast with one study, which used PCR to detect recKoRV using primers that spanned the PhER region of recKoRV and the 3’ KoRV LTR. In that study recKoRV was not detected in any koalas sampled from the Mornington Peninsula (n = 5) or Gippsland (n = 11) in Victoria (Löber *et al*. 2018), the PCR primers used in that study (Ya_recKoRV1-F, Ya_recKoRV1-R, recKoRV-F1, and KoR27-R), were compared in silico to the assembled recKoRV fragments utilised in this study for primer design. Binding sites for these primers, with 100% identity, were present in both Victorian koala recKoRVs assembled, one of which originated from the Strzelecki region of Gippsland. However notably of the assembled fragments, several from a South Australian animal lacked the forward primer binding region from (Löber *et al*. 2018). It may be that our sample set is simply missing recKoRV negatives from Victorian koala populations, but based on our findings, and those in South Australia (Stephenson *et al*. 2021), it is more probable that Southern koalas detected free of recKoRV are a result of issues with diagnostic sensitivity. Any subsequent Victorian or South Australian koalas thought to be negative for recKoRV should be further invested using advanced methods such as CRISPR based enrichment and long read sequencing, which may provide the most robust evidence for absence of integrated KoRV (Tarlinton *et al*. 2022).

A recent study showed non-KoRV-A subtypes emerged only when individuals had a sufficient amount of endogenised KoRV-A (at least one copy per cell) (Sarker *et al*. 2021; Blyton *et al*. 2022). If the presence of recKoRV in turn reduces the ability for KoRV to spread in a host, then this mechanism may inhibit significant KoRV associated disease by hampering the opportunity for non-KoRV-A subtypes to emerge within the individuals. Indeed, an extremely low KoRV subtype diversity was detected previously in Victorian populations with no or few KoRV-B (Legione *et al*. 2017; Sarker *et al*. 2021). Given KoRV-B is often more strongly associated with neoplasia (Quigley *et al*. 2018) and chlamydia (Waugh *et al*. 2017), it is possible that the high recKoRV copy numbers in Victorian populations might reduce these risks. Nevertheless, neoplasia in koalas is most likely a result of altered host genome transcription by the effect of KoRV LTRs (Hobbs *et al*. 2017) as in similar retroviruses such as Murine Leukemia Virus (Hanecak *et al*. 1991). As recKoRV copies still possess these transcriptionally active LTRs, their presence itself might still be pathogenic. On that account, our assay, which can examine the combined amount of intact KoRV and recKoRV, may also hold a significance as a diagnostic tool, to indicate individual animal susceptibility to KoRV-induced neoplasia.

Our findings have significant implications for management of koalas in Victorian populations. Current population management strategies and national risk assessments have included KoRV as an important disease, and the concept of remaining free of KoRV considered a possibility.

However, this research identifies that endogenous recKoRV is likely present in all koala populations and future research should focus on the role of recKoRV in exogenous KoRV and disease presentation in koalas.

## Acknowledgements

The authors wish to acknowledge the various contributors of the archived diagnostic samples that led to this research being possible.

