## Supplementary Table 01 for "A High Frequency of Detection of Recombinant Koala Retrovirus (recKoRV) in Victorian Koalas Suggests Historic Integration of KoRV"

**Supplementary Table 1.** Comparisons of KoRV-5’ prevalence and associated statistical analysis results

| **Variable** | **KoRV-5’ positive/n** | **Prevalence (%)** | **Odds ratio** | **95% CI** | **P value** | **KoRV-*pol* positive / KoRV-5’ positive** |
| --- | --- | --- | --- | --- | --- | --- |
| Sex* |  |  |  |  |  |  |
| Male | 108/110 | 98.2 | 0.64 | (0.08, 5.42) | 0.66 | 24/108 (22.2%) |
| Female | 168/170 | 98.8 | 1.00 | - | - | 39/168 (23.2%) |
| Not recorded | 6/7 | 85.7 | - | - | - | 2/6 (33.3%) |
| Age* |  |  |  |  |  |  |
| Young | 54/54 | 100 | - | - | - | 15/54 (27.8%) |
| Mature | 194/197 | 98.5 | 1.00 | - | - | 39/194 (20.1%) |
| Old | 21/22 | 95.4 | 0.33 | (0.04, 6.72) | 0.339 | 6/21 (28.6%) |
| Not recorded | 13/14 | 92.9 | **-** | - | - | 5/13 (38.5%) |
| Region * ^+^ |  |  |  |  |  |  |
| Far West | 18/18 | 100 | **-** | - | - | 6/18 (33.3%) |
| South Coast | 143/145 | 98.6 | 1.00 | - | - | 23/143 (16.1%) |
| Far North | 15/15 | 100 | - | - | - | 6/15 (40%) |
| Mornington Peninsula | 15/15 | 100 | - | - | - | 4/15 (26.7%) |
| French Island | 17/17 | 100 | - | - | - | 1/17 (5.9%) |
| Gippsland | 32/33 | 97.0 | 0.45 | (0.042, 9.80) | 0.52 | 6/32 (18.8%) |
| Raymond Island | 34/35 | 97.1 | 0.48 | (0.044, 10.41) | 0.55 | 18/34 (52.9%) |
| Others/ not recorded | 8/9 | 88.9 | - | - | - | 1/8 (12.5%) |
| Extraction Plate^ |  |  |  |  |  |  |
| 1 | 76/78 | 97.4 | 0.48 | (0.022, 5.06) | 0.55 | 20/76 (26.3%) |
| 2 | 76/80 | 95.0 | 0.24 | (0.012, 1.65) | 0.20 | 14/76 (18.4%) |
| 3 | 78/80 | 97.5 | 0.49 | (0.022, 5.19) | 0.56 | 9/78 (11.5%) |
| 4 | 80/81 | 98.8 | 1.00 | - | - | 28/80 (35%) |
| Sample type^ |  |  |  |  |  |  |
| Buffy coat | 142/147 | 96.6 | 1.00 | - | - | 21/142 (14.8%) |
| Spleen | 125/128 | 97.7 | 1.47 | (0.35, 7.27) | 0.61 | 33/125 (26.4%) |
| Whole blood | 34/34 | 100 | - | - | - | 17/34 (50%) |
| Plasma | 4/5 | 80 | 0.14 | (0.016, 3.02) | 0.10 | 0/4 (0%) |
| Serum | 5/5 | 100 | - | - | - | 0/5 (0%) |

* Each individual animal represented once. ^ Individual animals may be represented by multiple sample types. + Regions listed from west to east geographically.

CI: confidence interval. –: not measure
